# *Salmonella* Outer Membrane Vesicles contain tRNA Fragments (tRFs) that Inhibit Bacteriophage P22 infection

**DOI:** 10.1101/2021.11.09.467952

**Authors:** Dominika Houserova, Yulong Huang, Mohan V. Kasukurthi, Brianna C. Watters, Fiza F. Khan, Raj V. Mehta, Neil Y. Chaudhary, Justin T. Roberts, Jeffrey D. DeMeis, Trevor K. Hobbs, Kanesha R. Ghee, Cameron H. McInnis, Nolan P. Johns, Abrianna J. Kegler, Alexander B. Coley, Cana L. Brown, Jenny L. Hewes, Marie M. McElyea, Monica N. Reeves, Tuan M. Tran, Natalie R. Bauer, Jingshan Huang, Jonathon P. Audia, John W. Foster, Glen M. Borchert

## Abstract

*Salmonella* Outer Membrane Vesicles (OMVs) were recently shown to inhibit P22 bacteriophage infection. Furthermore, despite there being several published reports now independently describing (1) the marked prevalence of tRFs within secreted vesicle transcriptomes and (2) roles for specific tRFs in facilitating/inhibiting viral replication, there have been no examinations of the effects of vesicle-secreted tRFs on viral infection reported to date. Notably, while specific tRFs have been reported in a number of bacteria, the tRFs expressed by salmonellae have not been previously characterized. As such, we recently screened small RNA-seq datasets for the presence of recurrent, specifically excised tRFs and identified 31 recurrent, relatively abundant tRFs expressed by *Salmonella enterica* serovar Typhimurium (SL1344). What’s more, we find *S*. Typhimurium OMVs contain significant levels of tRFs highly complementary to known *Salmonella enterica*-infecting bacteriophage with 17 of 31 tRFs bearing marked complementarity to at least one known *Salmonella enterica*-infecting phage (averaging 97.4% complementarity over 22.9 nt). Most notably, tRNA-Thr-CGT-1-1, 44-73, bears 100% sequence complementary over its entire 30 nt length to 29 distinct, annotated *Salmonella enterica*-infecting bacteriophage including P22. Importantly, we find inhibiting this tRF in secreted OMVs improves P22 infectivity in a dose dependent manner whereas raising OMV tRF levels conversely inhibits P22 infectivity. Furthermore, we find P22 phage pre-incubation with OMVs isolated from naïve, control SL1344 *S*. Typhimurium, successfully rescues the ability of *S*. Typhimurium transformed with a specific tRNA-Thr-CGT-1-1, 44-73 tRF inhibitor to defend against P22. Collectively, these experiments confirm tRFs secreted in *S*. Typhimurium OMVs are directly involved with and required for the ability of OMVs to defend against bacteriophage predation. As we find the majority of OMV tRFs are highly complementary to an array of known *Salmonella enterica*-infecting bacteriophage, we suggest OMV tRFs may primarily function as a broadly acting, previously uncharacterized innate antiviral defense.

## INTRODUCTION

Developmentally regulated cleavage of tRNAs was initially reported in *Streptomyces coelicolor* in 2008^1^. In 2009, however, tRNA-derived RNA fragments (tRFs) of ~20 to 30 nt in length were first recognized as a class of functional small non-coding RNAs by a number of laboratories^2–4^. Since these initial reports, tRFs have been identified in all domains of life^5^. Despite this, how tRFs are generated and the functional roles of the majority of tRFs remain unclear although tRFs have been suggested to have initially arisen as a part of an ancient viral defense^6^. Supporting this concept, human endogenous retrovirus transposition rates can be limited by host tRF expression^7^, and specific tRFs are induced by viral infections (e.g., by RSV^8,9^). Notably, increasing titers of HIV have been shown to trigger the production of lysine tRFs that inhibit HIV, and T cells infected with human T-cell leukemia virus type 1 (HTLV-1) similarly upregulate the production of proline tRFs targeting this virus^10^. Together, these findings suggest that tRFs may constitute an uncharacterized component of the innate antiviral response.

Notably, tRFs have recently been found to be significantly enriched in Outer Membrane Vesicles (OMVs)^11^. OMVs are membrane-encapsulated spherical structures ~50 to 250 nm in diameter derived from Gram-negative bacteria cell envelopes^12^. OMVs are primarily generated by outer membrane blebbing but contain proteins, DNA, and RNAs at concentrations distinct from that of the intracellular complement^11,12^. OMVs have been associated with a number of different cellular functions including detoxification, pathogenicity, intercellular communication, and resistance to stressors^13^. Importantly, in 2015, Ghosal et al. found the majority of RNA contained within *Escherichia coli* OMVs has a length < 60 nt with an enrichment between 15 and 40 nts, and that *E. coli* OMV RNA sequencing reads primarily correspond to tRFs^10^. More recently, Koeppen et al.^14^ likewise found tRFs levels significantly higher in RNA isolated from *Pseudomonas aeruginosa* OMV than in intracellular RNA suggesting tRFs are actively enriched in OMVs prior to secretion. Interestingly, despite tRNAs and tRFs typically constituting only ~15% and 2% of normal intracellular small RNA populations respectively, tRFs have now been reported by several groups as being similarly abundant (representing >20% of RNA compositions) in secreted exosomal vesicles isolated from human saliva^15^, plasma^16^, semen^17^, mast cells^17^, mesenchymal stem cells^18^, and neurons^19^.

In summary, despite there being several published reports now independently describing (1) roles for specific tRFs in facilitating/inhibiting viral replication^3,8–10,20^, (2) the marked prevalence of tRFs within secreted vesicles (both exosomes and OMVs)^11,14–18^, and (3) the ability of OMVs to inhibit phage infection of several bacterial species (e.g., *E. coli*^21^, *Prochlorococcus marinus*^22^, *Shigella flexneri*^23^, *Salmonella enterica*^24^, and *Vibrio cholerae*^25^) there have been no examinations of the effects of vesicle-secreted tRFs on viral infection reported to date. Therefore, in light of a study in 2020 that conclusively demonstrated the ability of *Salmonella enterica* serovar Typhimurium OMVs to defend against bacteriophage P22^24^, we recently elected to define the full repertoire of *Salmonella enterica* serovar Typhimurium tRFs and explore their potential involvement in combating bacteriophage predation.

## RESULTS

### *Salmonella enterica* serovar Typhimurium tRFs

Specific tRFs have been reported in a number of bacteria^11,26^. To date, however, the tRFs expressed by *S*. Typhimurium have not been characterized. As such, we recently screened 12 preexisting small RNA seq datasets: six generated by our group^27^ and six by others^28,29^ for the presence of recurrent, specifically excised tRFs. Through employing a novel RNA-seq analysis platform designed to annotate fragments excised from longer noncoding RNAs, (SURFr)^30,31^, we have successfully identified 31 recurrent, relatively abundant tRFs expressed by *S*. Typhimurium. These tRFs average 26.7 nt in length ranging between 19 nt for the shortest (tRNA-Pro-TGG-1-1, 49-75) and 38 nt for the longest (tRNA-fMet-CAT-2-1, 33-70). All but four tRFs were found to be expressed under multiple conditions, and the median maximum tRF expression for all tRFs was 1,799 RPM with a mean of 4,333 RPM (**Table 1**). Notably, we find tRFs are similarly abundant in *E. coli, V. cholera, Staphylococcus aureus*, and *P.aeruginosa* small RNA transcriptomes (**Supplemental Tables 1-4**).

**Table 1.**
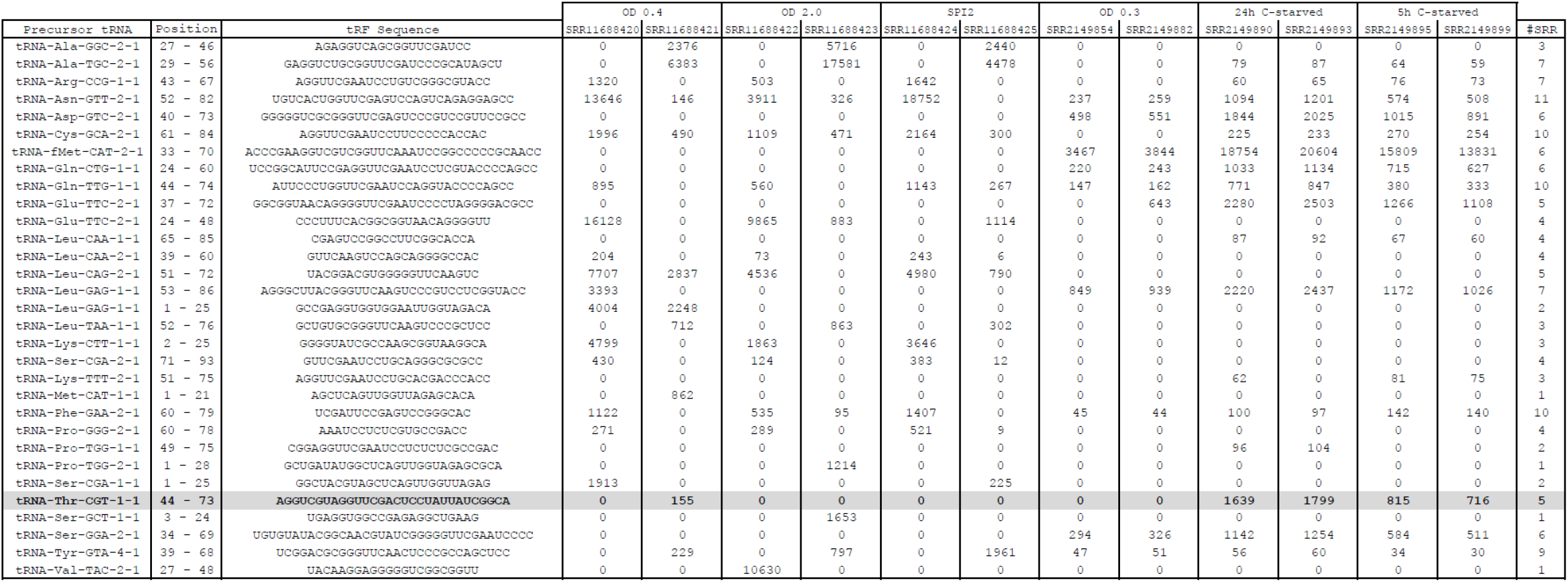
31 tRFs expressed in *S*. Typhimurium SL1344. SURFr-identified tRF precursor positions and excised sequences are shown. OD 0.4, Mid-log phase. OD 2.0, Stationary-phase. SPI2, salmonella pathogenicity island 2 conditions: (SPI-2 medium, OD600 = 0.3)^28,29^. OD 0.3, 24h and 5h C-starved from ^27^.

### tRNA-Thr-CGT-1-1, 44-73

In order to confirm the identities of SURFr-called tRFs, we selected tRNA-Thr-CGT-1-1, 44-73 for experimental validation. SURFr reported a 30 nt tRF specifically excised from the 3’ end of tRNA-Thr-CGT-1-1 at positions 44-73 (**Figure 1A-C**). While we find little expression of this tRF during exponential growth, we find it is robustly expressed (avg. 1,025 RPM) during Carbon starvation (**Table 1**). Importantly, subsequent qRT-PCRs confirm the expression of this specific tRF in response to Carbon starvation (**Figure 1D**).

**Figure 1.**
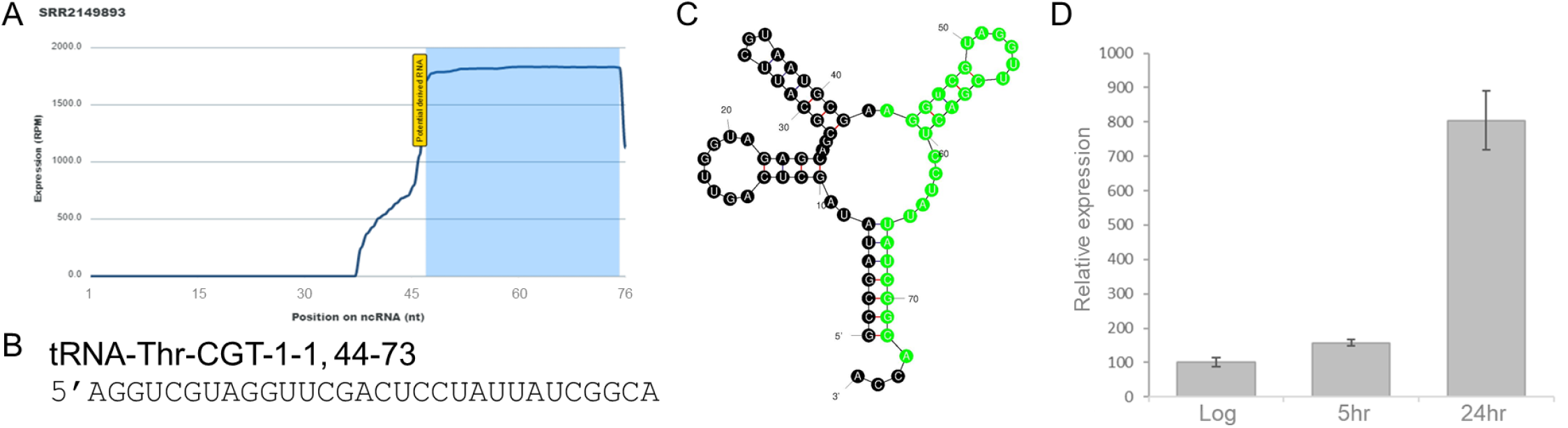
tRNA-Thr-CGT-1-1, 44-73. (A) SURFr^30,31^ “Differential Expression Vector” window depicting each nucleotide within tRNA-Thr-CGT-1-1 and indicating the fragment recurrently identified with a blue rectangle. The x-axis represents the position in the ncRNA selected (tRNA-Thr-CGT-1-1), and the y-axis depicts the expression level of the ncRNA at each position. (B) tRNA-Thr-CGT-1-1, 44-73 tRF sequence corresponding to the blue rectangle in “A”. (C) Secondary structure of tRNA-Thr-CGT-1-1 with positions 44-73 highlighted in green. Structural prediction generated with mFold^48^. (D) qRT-PCR confirming the increased expression of tRNA-Thr-CGT-1-1, 44-73 tRF in *S*. Typhimurium during carbon starvation (n=3). Relative expression, % expression as compared to average Log expression. Log, logarithmic phase. 5h, 5 hour carbon starved. 24h, 24 hour carbon starved.

### *S*. Typhimurium tRFs are highly complementary to bacteriophage

As tRNA fragments were first observed during bacterial phage infections^32^, we next evaluated the potential for these tRFs to interact with bacteriophage through direct basepairing. Strikingly, we found 17 of 31 SURFr-called *S*. Typhimurium tRFs bear marked complementarity to at least one known *S*. Typhimurium phage with an average sequence alignment of 97.4% complementarity over 22.9 nt. Of note, only two of these seventeen tRFs were identified as being complementary to a single bacteriophage. The other fifteen tRFs averaged significant complementarities to multiple, distinct phage (avg. 7.1) with tRNA-Leu-CAA-2-1, 39-60 significantly aligning to 24 phage, tRNA-Lys-TTT-2-1, 51–75 significantly aligning to 32 phage, and tRNA-Thr-CGT-1-1, 44-73 significantly aligning to 29 phage (**Supplemental Table 5**). In all, significant complementarities to 107 of 475 *Salmonella enterica*-infecting bacteriophage genomes available at the bacterial bioinformatics PathoSystems Resource Integration Center (PATRIC)^33^ were identified. Perhaps most notably, the tRF detailed in **Figure 1**, tRNA-Thr-CGT-1-1, 44-73, was found to be 100% complementary over its entire 30 nt length to 29 distinct annotated *Salmonella enterica*-infecting bacteriophage genomes (**Supplemental Table 6**).

### tRNA-Thr-CGT-1-1, 44-73 tRF sequestration improves P22 infectivity

In light of the striking complementarities between *Salmonella enterica*-infecting phage and SURFr-identified tRFs, we next asked if tRNA-Thr-CGT-1-1, 44-73 tRF expression was inhibitory to bacteriophage infection. We selected P22 for our analyses as tRNA-Thr-CGT-1-1, 44-73 tRF bears 100% sequence complementary over its entire 30 nt length to P22 bacteriophage (**Figure 2A**), and since P22 is extremely well characterized and has long been an important tool for investigating salmonella genetics^34^. We began by determining the effects of sequestering tRFs intracellularly. To achieve this, we preemptively transformed *S*. Typhimurium with antagomiR oligonucleotides perfectly complementary to tRNA-Thr-CGT-1-1, 44-73 tRF (Anti), identical to this tRF (Pos), or scrambled controls then assessed P22 infectivity via plating efficiency. We find preemptive transformation of *S*. Typhimurium with an antagomiR oligonucleotide designed to specifically bind and inhibit tRNA-Thr-CGT-1-1, 44-73 tRF (Anti) improved P22 infectivity in a dose dependent manner with antagomiR transformation improving P22 infectivity by ~60% at a 10^−3^ P22 dilution and nearly 450% at a 10^−4^ dilution (**Figure 2B,C**). In contrast, transformation with control antagomiRs with sequences either unrelated (scrambled) or identical to this tRF (Pos) did not significantly affect infectivity. Importantly, plating efficiency was assessed in unstarved cells as we find relatively limited expression of this tRF during growth in carbon-containing media (**Table 1**).

**Figure 2.**
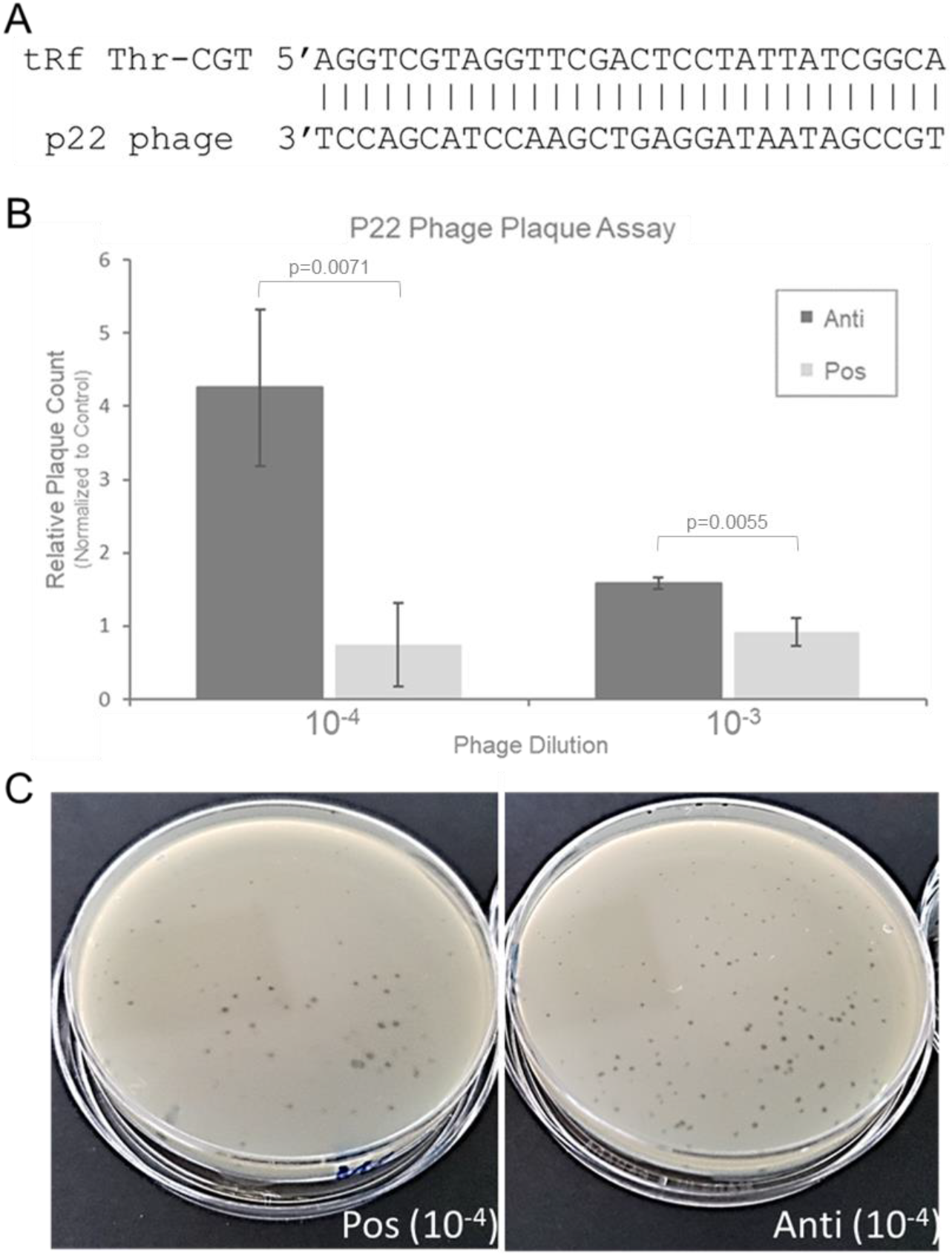
tRNA-Thr-CGT-1-1, 44-73 sequestration enhances P22 infectivity. (A) tRNA-Thr-CGT-1-1, 44-73 tRF alignment with P22 bacteriophage. (B) P22 plaque assay following preemptive transformation of *S*. Typhimurium with an antagomiR oligonucleotide (Anti) designed to specifically bind and inhibit tRNA-Thr-CGT-1-1, 44-73 tRF or its reverse complement (Pos). Plaque count was normalized to control: *S*. Typhimurium transfected with scrambled antagomiR oligonucleotide (Scram). Bacteria were transformed (CaCl2 heat shock) with antagomiR or control oligo and allowed to recover then viral lysate was added (at the indicated dilutions), mixed with molten top agar, and poured onto LB agar plates. Once solidified, plates were incubated at 37°C for 24hrs then plaques were enumerated. (n=3, p-values determined by unpaired t-test). (C) Representative P22 (10^−4^ dilution) plaque assay.

### *S*. Typhimurium OMV inhibition of P22 requires tRNA-Thr-CGT-1-1, 44-73 tRF

*S*. Typhimurium OMVs were recently reported to inhibit bacteriophage P22 infection^24^. Therefore, in light of having found that tRNA-Thr-CGT-1-1, 44-73 tRF intracellular sequestration improves P22 infectivity, we next explored the potential role of tRNA-Thr-CGT-1-1, 44-73 tRF in OMV-mediated P22 restriction. SURFr^30,31^ analysis of available *S*. Typhimurium OMV RNA-sequencing data (SRR6843039)^35^ and qRT-PCR of our own OMV RNA isolates both confirm the presence of tRNA-Thr-CGT-1-1, 44-73 tRF within *S*. Typhimurium OMVs. What’s more, we find tRNA-Thr-CGT-1-1, 44-73 tRF levels are ~3.5x higher in OMV RNA isolates than in total cellular RNA (**Figure 3A**). Notably, the average diameter of our isolated vesicles (119.1 nm) strongly agrees with OMV identity^11,12^ (**Figure 3B**), and fluorescence-activated cell sorting following Texas Red labeled antagomiR internalization confirms OMV isolates are readily transfectable (**Figure 3C**).

**Figure 3.**
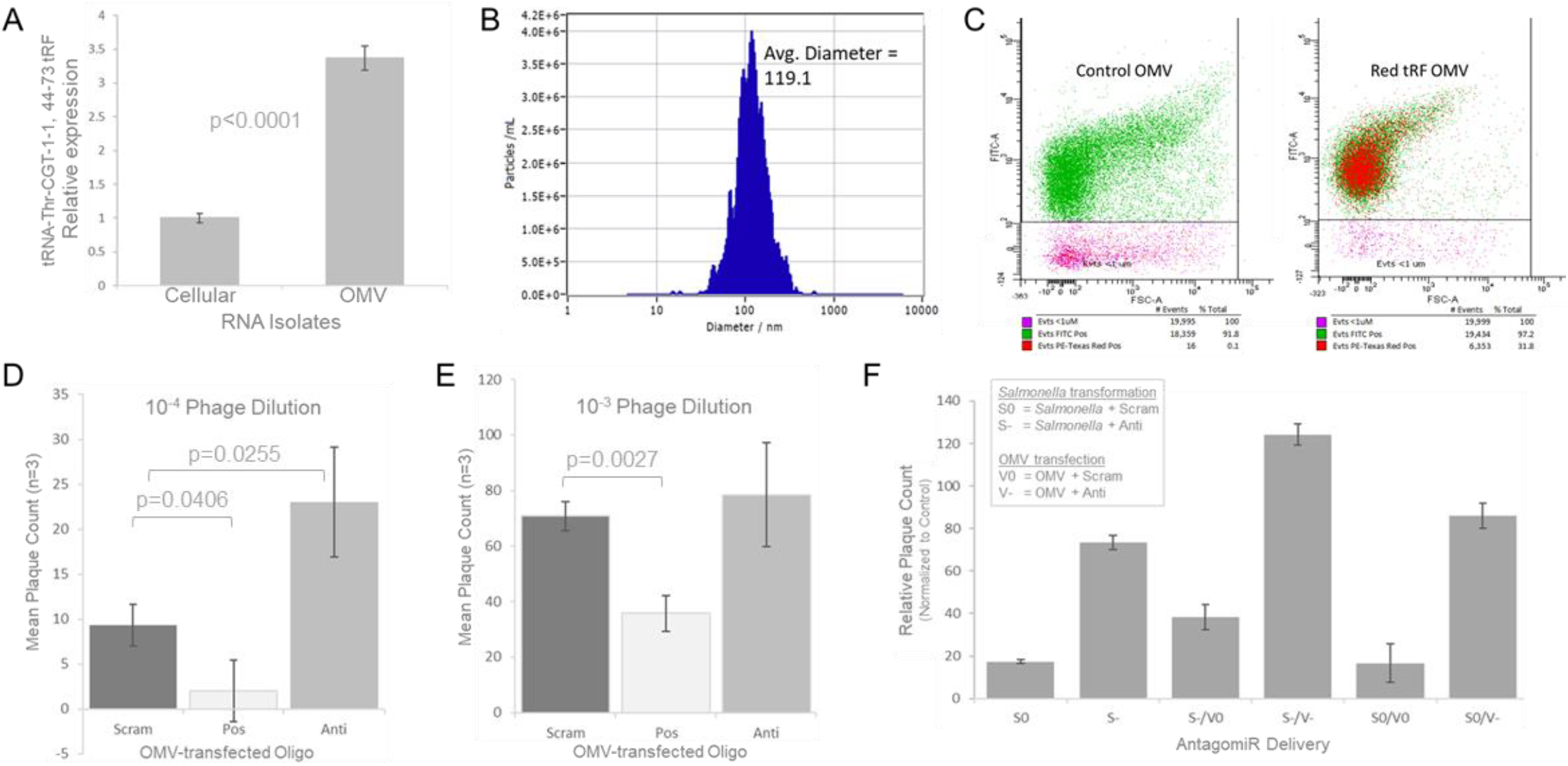
*S*. Typhimurium OMV inhibition of P22 requires tRNA-Thr-CGT-1-1, 44-73 tRF. (A) qRT-PCR confirming the expression of tRNA-Thr-CGT-1-1, 44-73 tRF in *S*. Typhimurium OMV RNA. Relative expression, % expression as compared to average total cellular RNA expression (n=3, p-value determined by unpaired t-test). Cellular, total cellular RNA isolate. OMV, OMV RNA isolate. (B) *S*. Typhimurium OMVs were harvested then analyzed at a 1:100 dilution on ZetaView. Concentration: 1.3E8 particles / mL. (C) 1×10^6^ OMVs from each sample were transfected with 1μM scrambled control oligo (left) or 1μM Texas Red labeled antagomiR (right) then analyzed by Aria FACS. Dot plots show fluorescent intensities of 10,000 events; maximal acquisition time was 6 sec and sample pressure was < 3. Percentages of gated populations are shown. (D) Plaque Assay (10^−4^ dilution) following P22 phage pre-incubation with OMVs transfected with 1μM scrambled control (Scram), an antagomiR oligo designed to specifically bind and inhibit tRNA-Thr-CGT-1-1, 44-73 tRF (Anti), or its reverse complement (Pos). (n=3, p-value determined by unpaired t-test). (E) Plaque Assay (10^−3^ dilution), otherwise as in “D”. (F) Plaque assay (10^−4^ dilution). “S0”, *S*. Typhimurium transformed with Scram antagomiR control / no phage preincubation with OMVs. “S-”, *S*. Typhimurium transformed with Anti antagomiR / no phage preincubation with OMVs. “S-/V0”, *S*. Typhimurium transformed with Anti antagomiR / phage preincubated with OMVs transfected with Scram antagomiR control. “S-/V-”, *S*. Typhimurium transformed with Anti antagomiR / phage preincubated with OMVs transfected with Anti antagomiR. “S0/V0”, *S*. Typhimurium transformed with Scram antagomiR control / phage preincubated with OMVs transfected with Scram antagomiR control. “S0/V-”, *S*. Typhimurium transformed with Scram antagomiR control / phage preincubated with OMVs transfected with Anti antagomiR.

As such, we proceeded by preincubating P22 phage with OMVs transfected with antagomiR oligonucleotides perfectly complementary to tRNA-Thr-CGT-1-1, 44-73 tRF (Anti), identical to this tRF (Pos), or scrambled control then assessed the effects of preincubation with OMVs on P22 infectivity by measuring plating efficiency in naïve, untransformed *S*. Typhimurium. Excitingly, we find P22 phage pre-incubation with OMVs containing the antagomiR shown to inhibit tRNA-Thr-CGT-1-1, 44-73 tRF and improve P22 infectivity in a dose dependent manner following transformation (**Figure 2**) similarly improved P22 infectivity of naïve control *S*. Typhimurium by ~250% at a 10^−4^ P22 dilution whereas preincubation with OMVs containing elevated tRNA-Thr-CGT-1-1, 44-73 tRF levels (Pos) conversely inhibited P22 infectivity by >75% (**Figure 3D,E**) as compared to controls. Importantly, we find P22 phage pre-incubation with OMVs isolated from naïve control *S*. Typhimurium successfully rescues the ability of *S*. Typhimurium transformed with the tRNA-Thr-CGT-1-1, 44-73 tRF antagomiR to defend against P22 which strongly suggests OMV tRFs are primarily responsible for phage inhibition (**Figure 3F**). Finally, infectivity was not significantly altered following P22 pre-incubation with OMVs transfected with control antagomiRs as compared to infectivity following pre-incubation with naïve OMVs.

## DISCUSSION

For decades, it was widely believed that the sole role of tRNAs was to convert the information encoded on mRNAs into amino acid sequence. Despite the existence of tRNA-derived fragments (tRFs) being known since the 1990s, until quite recently they have generally been disregarded as degradation products. Notably, pieces of tRNAs were first observed during phage infections of bacteria^1^, and in humans, endogenous retrovirus transposition rates can be limited by host tRF expression and specific tRFs are induced by viral infection^3,8–10,20^. Together, these findings potentially suggest a role for tRFs in innate antiviral defense. However, while specific tRFs have been reported in a number of bacteria, neither (1) the tRFs expressed by salmonellae nor (2) roles for tRFs in combatting bacteriophage have been reported to date.

While specific tRFs have been reported in a number of bacteria, the tRFs expressed by salmonellae have not been previously characterized. As such, we recently screened small RNA-seq datasets for the presence of recurrent, specifically excised tRFs and identified 31 recurrent, relatively abundant tRFs expressed by *S*. Typhimurium. What’s more, *S*. Typhimurium OMVs can inhibit P22 bacteriophage infection^24^, and we find *S*. Typhimurium OMVs contain significant levels of tRFs highly complementary to known *Salmonella enterica*-infecting bacteriophage. Most notably, tRNA-Thr-CGT-1-1, 44-73, is 100% complementary to 29 distinct, annotated *Salmonella enterica*-infecting bacteriophage including P22, and this tRF is 100% conserved in *E. coli* (**Supplemental Table 1**).Excitingly, we find inhibiting this tRF in secreted OMVs improves P22 infectivity in a dose dependent manner whereas raising OMV tRF levels conversely inhibits P22 infectivity. Furthermore, we find P22 phage pre-incubation with OMVs isolated from naïve, control *S*. Typhimurium, (and therefore containing unaltered, baseline tRF levels) successfully rescues the ability of *S*. Typhimurium preemptively depleted of tRNA-Thr-CGT-1-1, 44-73 tRF to defend against P22. Collectively, these experiments confirm tRFs secreted in *S*. Typhimurium OMVs are directly involved with and required for the ability of OMVs to defend against bacteriophage predation.

In agreement with the proposed role for OMV tRFs in combatting phage infection, the ability of OMVs to inhibit phage infection of several bacterial species (e.g. *E. coli*^21^, *P. marinus*^22^, *S. flexneri*^23^, *S. enterica*^24^, and *V. cholerae*^25^) has now been reported. What’s more, in addition to the confirmation that *S*. Typhimurium OMVs contain significant levels of tRFs reported here, tRFs have been reported to be significantly enriched in *E. coli* and *P. aeruginosa* OMVs. In 2015, Ghosal et al. found the majority of RNA contained within *E. coli* OMVs has a length < 60 nt with an enrichment between 15 and 40 nts, and that *E. coli* OMV RNA sequencing reads primarily correspond to tRFs. More recently, Koeppen et al.^14^ likewise found tRFs levels significantly higher in RNA isolated from *P. aeruginosa* OMVs than in intracellular RNA suggesting tRFs are actively enriched in OMVs prior to secretion.

Of note, in 2014, Biller et al. mixed purified *Prochlorococcus* vesicles with an infective phage (PHM-2) and used electron microscopy to record numerous examples of direct vesicle binding by phage. They found, many vesicle-attached phage had altered capsid staining density and a shortened stalk suggesting that they had injected their DNA into the vesicle. In light of this, they suggested their findings support a decoy model in which the export of vesicles by marine bacteria acts to reduce the likelihood of cellular infection through effectively reducing the number of phage available to infect cells^22^. Of note, our work contradicts elements of this model in that although cells are constantly releasing OMVs, we find the inhibition of specific OMV tRFs can significantly reduce the ability of OMVs to restrict phage infectivity suggesting the requirement for a direct interaction between tRFs and the phage they inhibit.

Also of interest, transfer RNA genes have long been known to represent the preferred chromosomal sites for bacteria prophage integration. Generally, both Gram-negative and Gram-positive prophages integrate into tRNA loci in the proper orientation for the phage *attP* site to reconstitute the tRNA upon entry^36^. Notably, the primary tRF examined in this work, tRNA-Thr-CGT-1-1, 44-73, bears 100% sequence complementary over its entire 30 nt length to 29 distinct, annotated *Salmonella enterica*-infecting bacteriophage including P22 (**Table 2**), and interestingly, the tRNA locus expressing this tRF also serves as the specific genomic site targeted by P22 for prophage integration^37,38^.

That said, how or if this tRF directly interacts with its complementary sequence at the *attP* site of the P22 genome remains unclear. Additional studies will clearly be required to determine whether this interaction occurs ex vivo and to specifically define the mechanism of tRF-based phage restriction including the extent of tRF sequence complementary necessary to confer regulation. While we found 17 of 31 SURFr-called *S*. Typhimurium tRFs bear marked complementarity to at least one known *Salmonella enterica*-infecting phage with an average sequence alignment of 97.4% complementarity over 22.9 nt to date, we have only definitively confirmed the ability of tRNA-Thr-CGT-1-1, 44-73 (which bears 100% sequence complementary over its entire 30 nt length to P22) to restrict P22 phage infection. More fundamentally, the full repertoire of bacterial tRFs being expressed from salmonellae (and those expressed from virtually all other prokaryotes) as well as the tRFs specifically enriched in OMVs remain significant gaps in knowledge. While the 31 tRFs reported here represent the most comprehensive characterization of *S*. Typhimurium tRFs reported to date, we find the tRFs expressed under the six distinct growth conditions evaluated differ significantly suggesting further tRFs expressed during specific conditions and/or challenges not examined in the current study will likely be described. Similarly, we suggest the tRFs we identify in *E. coli* (54 tRFs from 82 SRR files), *V. cholera* (20 tRFs from 10 SRR files), *S. aureus* (31 tRFs from 7 SRR files), and *P. aeruginosa* (48 tRFs from 9 SRR files) small RNA transcriptomes (**Supplemental Tables 1-4** respectively) likely also represent significant, but incomplete, tRF catalogs.

Finally of note, tRFs unquestionably participate in more than viral defense. In eukaryotes, an array of regulatory roles for tRFs have been reported involving: transcription, translation, apoptosis, cellular proliferation / differentiation, RNAi, intercellular communication, epigenetics, retrotransposon restriction, etc^39^. While several bacterial tRF characterizations have also been reported^40–42^, to date, the functions tRFs serve in prokaryotes are far less well described. Notably, during alkaline stress, *Haloferax volcanii* has been shown to express a 5’ tRF that binds its small ribosomal subunit to interfere with mRNA loading and universally downregulate protein expression^43^. Also of note, bacterial tRFs have been reported to be involved in interspecies communication. Rhizobial tRFs were recently shown to be absorbed by symbiotic hosts and silence the expression of host genes through interacting with their RNA interference machinery^44^.

The work presented here clearly demonstrates that tRFs secreted in *S*. Typhimurium OMVs are directly involved with and required for the ability of OMVs to defend against bacteriophage predation. As we find the majority of OMV tRFs are highly complementary to an array of known *Salmonella enterica*-infecting bacteriophage, we speculate that OMV tRFs may primarily function as a broadly acting, previously uncharacterized ancient antiviral defense. Finally, as several published reports have now independently described (1) the marked prevalence of tRFs within exosomal transcriptomes^11,14–18^ and (2) roles for specific human tRFs in inhibiting viral replication^3,8–10,20^, it is tempting to also speculate that vesicle-secreted tRFs may similarly function as broad-spectrum antivirals in eukaryotes.

## METHODS

Unless stated otherwise all chemicals and kits were purchased from ThermoFisher. *Salmonella enterica* serovars Typhimurium strains (SL1344 and UK1) and aliquot of P22 HT105/1-int bacteriophage were provided by Dr. John Foster.

### SURFr tRF annotation and identification of phage complementarities

SRA IDs corresponding to bacterial small RNA-seq datasets were obtained from the NCBI SRA database^45^ and entered directly into SURFr, a real-time NGS data analytic tool to identify and analyze ncRNA-derived RNAs, for analysis after selecting the desired bacterial species^30,31^. Filter was set to “tRNA” only, and all tables, reports, and images downloaded. The SURFr platform is publically available at: http://salts.soc.southalabama.edu/surfr

Final SURFr-identified tRFs were aligned to all 475 *Salmonella enterica*-infecting bacteriophage genomes available through the bacterial bioinformatics PathoSystems Resource Integration Center (PATRIC)^33^ funded by the National Institute of Allergy and Infectious Diseases (https://www.patricbrc.org) using BLAST+ (2.2.27). All resulting alignments were filtered by requiring a minimum length of 17 base pairs with 100% sequence identity or 19+ bps with an evalue of 0.01 or less.

### Cultivation of bacterial cultures

Cultures of *S*. Typhimurium deprived of carbon source and control non-starved were generated as previously described^27^. Briefly, overnight cultures of SL1344 were inoculated into MOPS minimal media (MS) with either low (0.03%; loC) or high (0.4%; hiC) glucose content ^46^. Both cultures were then grown at 37°C with aeration/shaking until harvested; Non-starved cultures grown in MS-hiC were collected mid-log phase (OD600=0.4) while starved cultures grown in MS-loC were collected at 5 hours and 24 hours after reaching stationary phase to ensure complete depletion of C-source. For viral plaque assays and co-incubation experiments standard, commercially available Lysogeny Broth (LB) was used for all *S*. Typhimurium culturing.

### Validation of tRF presence via quantitative RT-PCR

Small RNA was isolated using mirVana miRNA Isolation Kit according to manufacturer’s instructions. Real-time, quantitative PCR was performed to validate presence of tRF-21 in UK1 strain of *S*. Typhimurium and its OMVs using All-in-One miRNA qRT-PCR Kit (GeneCopia). The reactions were performed in triplicate in a 96-well plates using 0.2 uM of each custom forward and universal reverse primers provided in the kit and 1.5 ug of total RNA in nuclease-free water. qRT-PCR was conducted on the iQ-5 Real-Time PCR Detection System (Bio-Rad) with following settings: initial polymerase activation and DNA denaturation at 95°C for 10 minutes, followed by 40 cycles of 95°C for 10 sec, 60°C for 20 sec, 72°C for 15 sec. Specificity of amplifications was verified using melting curves. Gene expression was calculated via the Delta-Delta cycle threshold method. qRT-PCR primers are listed in **Supplemental Table 7**.

### P22 bacteriophage propagation

An overnight culture of P22-susceptible *S*. Typhimurium strain UK1 was used to propagate the bacteriophage by overnight co-incubation at 37°C with constant shaking. The next day, bacteria were lysed by vigorous vortexing with 3 drops of chloroform and 100% ethanol. The phage was then precipitated from the lysate by addition of PEG6000 (final concentration 7% w/v) and NaCl (final concentration 0.5M). Solution was then incubated for 60 min at RT and phage was concentrated by centrifugation at 8000xg for 10 min. Phage pellet was then re-suspended in 100uL of 1X PBS and stored at 4°C.

### OMV isolation and validation

An overnight culture of *S*. Typhimurium strain UK1 was inoculated into standard minimal media that was depleted of iron (0.1% (NH4)2SO4; 0.9% KH2PO4, 0.2% glycerol, 0.04% MgSO4, with 0.5ug of Fe^2+^ per ml) and grown until late log/early stationary phase. The culture was then collected and cells and large cellular debris removed by centrifugation (5000xg for 15 mins) and two subsequent filtrations with 0.45 and 0.2 micron filters. OMVs were then isolated and concentrated using Pall Minimate Tangenial Flow Filtration system and 100 kDa filter cassette. Concentrated particles were further purified by ultracentrifugation (1hr, 120k x g, 4°C).The presence and quantity of OMVs was determined using ZetaView (Particle Metrix ZetaView system) and 1×10^6^ OMVs from each sample were transfected with 1μM scrambled control (scram) or 1μM Texas Red-labeled antagomiR (Anti-Red) (**Supplemental Table 7**) and stained using green fluorescent DiOC_18_(3) membrane dye. Samples were then analyzed and sorted on a Becton Dickinson FACS Aria II Analyzer and Cell Sorter. Transfection efficiency was calculated using number of OMVs positive for both green and red labels.

### tRF transformation and delivery via OMVs

*Intracellular tRF transformation*: An overnight culture of UK-1 was inoculated 1:100 into fresh LB broth media and allowed to grow until early to mid-log phase (OD=0.3). Once grown, the culture was chilled on ice for 10 minutes and the bacteria subsequently pelleted by centrifugation (3000X g, 10 minutes, 4°C). The supernatant was discarded and bacteria re-suspended in sterile, ice-cold 0.1M CaCl_2_ solution. The culture was then incubated on ice with shaking for additional 30 minutes. Bacteria were pelleted as before, re-suspended in sterile, ice-cold 15% glycerol in 0.1M CaCl_2_, aliquoted into 50μL aliquots and put on ice. 10μL of 100μM of tRF or control oligo was added to the aliquots and left undisturbed on ice for additional 30 minutes. The transformation was performed by heat shock at a 42°C water bath for 30 seconds. After transformation samples were incubated on ice for 2 minutes and subsequently diluted 1:1 in pre-warmed S.O.C media and incubated on a shaker at 37°C for 20 minutes. *OMV tRF transformation*: OMVs were transfected with either tRF mimic (Pos) or inhibitor (Anti) **Supplemental Table 7**. via electroporation using *E. coli* Pulser (Bio-Rad). Electroporation was performed under same conditions as previously described^47^. Transformed OMVs were put on ice and used immediately. *Viral Plaque Assay and Co-incubation*: To perform viral plaque assays, bacterial were grown and transfected with desired oligo as described above. After the last incubation, 10μL of viral lysate dilution was added and samples incubated at 37°C for additional 10 minutes. Samples were then transferred to a tube containing Molten Top Agar (1% Bacto tryptone, 0.5% Bacto yeast extract, 0.5% NaCl, 0.4% technical agar, 0.01M CaCl2, pH 7.4), mixed and poured onto prepared LB agar plates. Once solidified, plates were incubated at 37°C for 24hrs before the plaques were counted. For OMV tRF delivery experiments, *S*. Typhimurium were transformed with either tRF or control oligos and allowed to incubate in SOC (Super Optimal broth with Catabolite repression) as before. At the same time, 10 μL of viral lysate was added to 50 μL aliquots of OMVs-either naïve or transfected with tRF-21 oligo. Solutions were then placed on a shaking incubator (200 rpm) and allowed to incubate for 30 minutes. Transfected cells and OMV/viral solutions were then combined, and co-incubated additional 15 minutes before being plated in Molten Top Agar. Plates were incubated at 37°C for 24hrs and viral plaques were counted.

## DATA AVAILABILITY

All next-generation small RNA deep-sequencing libraries utilized are publicly available and were obtained from NCBI SRA. SRR Files analyzed: SRR6843039, SRR11688424, SRR2149890, SRR2149854, SRR2149895, SRR2149893, SRR11688421, SRR2149882, SRR11688423, SRR11688422, SRR11688425, SRR2149899, SRR11688420. All other relevant data (e.g. alignment files) are available from the authors upon request.

## ACKNOWLEDGEMENTS

We thank the University of South Alabama College of Medicine Department of Pharmacology for ongoing support. Funding was provided in part by NSF RAPID grant 2030080 (GMB) awarded by Division of Molecular and Cellular Biosciences (with co-funding provided by the NSF EPSCoR program), in part by NIH RO1HL133066 (NRB), and in part by the USA COM IGP grant 1828 (GMB). Graduate funding was also provided in part by Alabama Commission on Higher Education ALEPSCoR grants 150380 (JDD). The project used an instrument funded, in part, by the National Science Foundation MRI, grant number CNS-1726069.

## AUTHOR CONTRIBUTIONS

DH and GMB had full access to all of the data in the study and take responsibility for the integrity of the data and accuracy of the data analysis. This included study concept and design, experimental design and interpretation, data analysis, drafting of manuscript, critical revision of the manuscript for important intellectual content, and general study supervision. Each additional author contributed directly to the design of specific experiments, data analysis and to overall manuscript editing and approval.

## SUPPLEMENTAL TABLE LEGENDS

**Supplemental Table 1**. **54 tRFs expressed in *Escherichia coli*.**

**Supplemental Table 2. 20 tRFs expressed in *Vibrio cholera*.**

**Supplemental Table 3. 31 tRFs expressed in *Staphylococcus aureus*.**

**Supplemental Table 4. 48 tRFs expressed in *Pseudomonas aeruginosa*.**

**Supplemental Table 5. 17 *S*. Typhimurium tRFs are markedly complementary to bacteriophage sequences.** “Annotation” and “Start – End” refer to Ensembl tRNA-excised tRFs identified by SURFr^30,31^. Phage Name, correspond to phage genomes in the bacterial bioinformatics PathoSystems Resource Integration Center (PATRIC)^33^.

**Supplemental Table 6. tRNA-Thr-CGT-1-1, 44-73 is 100% complementary to 29 distinct, annotated *Salmonella enterica*-infecting bacteriophage genomes.** Accn, Common Name, Start and Stop positions all correspond to phage genomes in the bacterial bioinformatics PathoSystems Resource Integration Center (PATRIC)^33^.

**Supplemental Table 7. Oligonucleotide master list**. All oligonucleotides were synthesized by IDT DNA Technologies (Coralville, IA) at 100μM scale and HPLC purified.

