## Supplemental Table 5 for "*Salmonella* Outer Membrane Vesicles contain tRNA Fragments (tRFs) that Inhibit Bacteriophage P22 infection"

<table>
<tr><th>Annotation</th><th>Start - End</th><th>Phage Name</th><th>Alignment</th></tr>
<tr><td>tRNA-Asn-GTT-2-1</td><td>52 - 82</td><td>FSL SP-029, SE5, vB_SnwM_CGG4-1*</td><td>CACUGGUUCGAGUCCAGU<br>||||||||||||||||||<br>CACUGGUUCGAGUCCAGU</td></tr>
<tr><td>tRNA-Asp-GTC-2-1</td><td>40 - 73</td><td>38, BSP101, Mutine, PhiSH19, S8, S115, SE5, SE14, SeSz-1, SP1, SPAsTU*, vB_SalM_PM10, vB_SalM_SJ2</td><td>GGGGGUCGCGGGUUCGAGUCCCGUCCGUUCCGCC<br>|| |||||||||||||||||||||| || | |||<br>GGAGGUCGCGGGUUCGAGUCCCGUCAGUCCUGCC</td></tr>
<tr><td>tRNA-Gln-CTG-1-1</td><td>24 - 60</td><td>SE131*, SPN3US</td><td>CAUUCCGAGGUUCGAAUCCUCG<br>|| | |||||||||||||||||<br>CAGUACGAGGUUCGAAUCCUCG</td></tr>
<tr><td>tRNA-Gln-TTG-1-1</td><td>44 - 74</td><td>SPAsTU*</td><td>AUUCCCUGGUUCGAAUCCAGGUACCCCAGCC<br>|| ||||||||||||||||||| || | |||<br>AUGCCCUGGUUCGAAUCCAGGUGCCACUGCC</td></tr>
<tr><td>tRNA-Glu-TTC-2-1</td><td>37 - 72</td><td>FSL SP-010, Vi01*</td><td>GGGGUUCGAAUCCCCUA<br>|||||||||||||||||<br>GGGGUUCGAAUCCCCUA</td></tr>
<tr><td>tRNA-Leu-CAA-2-1</td><td>39 - 60</td><td>38, BSP101, GG32, Matapan, Melville, Mooltan, Mutine, PS5, S8, S117, S118, SenALZ1, SenASZ3, SeSz-1, SFP10, SJ46, SS3, SS9, STML-13-1, STML-198*, STP07, STP4-a, vB_SenM-2, Vi01</td><td>GUUCAAGUCCAGCAGGGGCCAC<br>||||||||||||||||||||||<br>GUUCAAGUCCAGCAGGGGCCAC</td></tr>
<tr><td>tRNA-Leu-GAG-1-1</td><td>1 - 25</td><td>SHP1*</td><td>GAGGUGGUGGAAUUGGUAGA<br>|||||||||||| |||||||<br>GAGGUGGUGGAAGUGGUAGA</td></tr>
<tr><td>tRNA-Lys-CTT-1-1</td><td>2 - 25</td><td>SPAsTU*</td><td>GGUAUCGCCAAGCGGUAAGGC<br>||||| |||||||||||||||<br>GGUAUAGCCAAGCGGUAAGGC</td></tr>
<tr><td>tRNA-Lys-TTT-2-1</td><td>51 - 75</td><td>7t3, Munch*, S113, S147, Stp1,</td><td>AGGUUCGAAUCCUGCACG<br>||||||||||||||||||<br>AGGUUCGAAUCCUGCACG</td></tr>
<tr><td>tRNA-Met-CAT-1-1</td><td>1 - 21</td><td>FSL SP-029, FSL SP-058, FSL SP-076, SPAsTU*</td><td>AGCUCAGUUGGUUAGAGC<br>||||||||||||||||||<br>AGCUCAGUUGGUUAGAGC</td></tr>
<tr><td>tRNA-Pro-TGG-2-1</td><td>1 - 28</td><td>41, SJ46*</td><td>GCUCAGUUGGUAGAGCGC<br>||||||||||||||||||<br>GCUCAGUUGGUAGAGCGC</td></tr>
<tr><td>tRNA-Ser-CGA-1-1</td><td>1 - 25</td><td>FSL SP-029, SPAsTU*</td><td>CGUAGCUCAGUUGGUUAGAG<br>||||||||||||||||||||<br>CGUAGCUCAGUUGGUUAGAG</td></tr>
<tr><td>tRNA-Lys-TTT-2-1</td><td>51 - 75</td><td>3-29, 19, 41, 100268_sal2, 118970_sal2, epsilon34, LVR16A, Meda, OSY-STA, S114, S124, S133, SE8, SE11(x2), Seabear, Seafire, SEN22*, SKML-39, SP01, SP1a, SP3, SPAsTU, SPC32H, SPC32N, SPN1S, SPN9TCW, STG2, Stitch, Sw2, Th1, vB_SenS_PHB06, vB_SenS_SB6</td><td>AGGUUCGGAAUCCUGCA<br>|||||||||||||||||<br>AGGUUCGGAAUCCUGCA</td></tr>
<tr><td>tRNA-Ser-CGA-2-1</td><td>71 - 93</td><td>epsilon34, SEN22*</td><td>GUUCGAAUCCUGCAGGGCGCGCC<br>|||||||||||||||||||||||<br>GUUCGAAUCCUGCAGGGCGCGCC</td></tr>
<tr><td>tRNA-Ser-GGA-2-1</td><td>34 - 69</td><td>7t3, Munch, PVP-SE1*</td><td>AUCGGGGGUUCGAAUCCC<br>||||||||||||||||||<br>AUCGGGGGUUCGAAUCCC</td></tr>
<tr><td>tRNA-Thr-CGT-1-1</td><td>44 - 73</td><td>22, 25, 34, 64795 sal4, 101962B sal5, 118970 sal4, 146851 sal5, 103203 sal4, 103203 sal5, 146851 sal4, Bp96115, FSL SP-010, GE vB TR, P22, S107, S135, S137, S149, SE10, SE16(x2), SE21, SE22, SF3, SF11, SI23, SE1 (in:p22virus)(x2), SPN9CC*, ST160, vB SalP PM43</td><td>AGGUCGUAGGUUCGACUCCUAUUAUCGGCA<br>||||||||||||||||||||||||||||||<br>AGGUCGUAGGUUCGACUCCUAUUAUCGGCA</td></tr>
<tr><td>tRNA-Tyr-GTA-4-1</td><td>39 - 68</td><td>RE-2010(x2), SEN8*</td><td>UCGGACGCGGGUUCAACUCCCGCCAGCUCC<br>||||||||||||||||||||||||||||||<br>UCGGACGCGGGUUCAACUCCCGCCAGCUCC</td></tr>
</table>

1.(x2): Alignments at 2 positions

2.(*): Indicates best alignment

3. The tRf extensively studied in this manuscript (tRNA-Thr-CGT-1-1, 44-73) is bolded.
