## Supplemental Table 7 for "*Salmonella* Outer Membrane Vesicles contain tRNA Fragments (tRFs) that Inhibit Bacteriophage P22 infection"

**Supplemental Table 7**. Oligonucleotide master list. All oligonucleotides were synthesized by IDT DNA Technologies (Coralville, IA) at 100µM scale and HPLC purified.

| Gene | Sequence (5’-3’) |
|---|---|
| tRNA-Thr-CGT-1-1, 44-73 tRF qRT-PCR | F: GGTCGTAGGTTCGACTCCTATTATCGG |
| | R: Qiagen All-in-one miRNA qRT-PCR Rev (#QP029) |
| STnc3920 qRT-PCR control | F: TAAAGCTCCCCGTAATTTAGCA |
| | R: CAGGGCGTTAATTACCTTTGAA |
| IsrL qRT-PCR control | F: CCGTTAACTGGCATCCTTCTAT |
| | R: GGCGACCTCTATTTGTTCATTC |
| tRNA-Thr-CGT-1-1, AntagomiR (Anti) | mG/ZEN/mCmCmGmAmUmAmAmUmAmGmGm<br>AmGmUmCmGmAmAmCmCmUmAmCmG/3ZEN/ |
| tRNA-Thr-CGT-1-1, Mimic (Pos) | mU/ZEN/mCmGmUmAmGmGmUmUmCmGmAm<br>CmUmCmCmUmAmUmUmAmUmCmGmG/3ZEN/ |
| tRNA-Thr-CGT-1-1, Scrambled Control (Scram) | mG/ZEN/mUmAmCmUmCmCmAmGmUmGmAm<br>CmAmAmAmAmGmCmCmGmUmAmGmG/3ZEN/ |
| tRNA-Thr-CGT-1-1, Texas Red (Anti-Red) | 5TEX615/mGmCmCmGmAmUmAmAmUmAm<br>GmGmAmGmUmCmGmAmAmCmCmUmAmCmG |
